# Lack of rhythmicity in Bmal1 deficient mice impairs motivation towards natural stimuli

**DOI:** 10.1101/2024.02.07.579260

**Authors:** Paula Berbegal-Sáez, Ines Gallego-Landin, Javier Macía, Laia Alegre-Zurano, Adriana Castro-Zavala, Patrick-Simon Welz, Salvador A Benitah, Olga Valverde

**Affiliations:** Neurobiology of Behavior Research Group (GReNeC-NeuroBio), Department of Medicine and Life Sciences (MELIS), Universitat Pompeu Fabra, Barcelona, Spain; Synthetic Biology for Biomedical Applications, Department of Medicine and Life Sciences (MELIS), Universitat Pompeu Fabra, Barcelona, Spain; Program in Cancer Research, Hospital del Mar Research Institute (IMIM), Barcelona, Spain; Institute for Research in Biomedicine (IRB Barcelona), Barcelona Institute of Science and Technology, 08028 Barcelona, Spain; Catalan Institution for Research and Advanced Studies (ICREA), Barcelona, Spain; Catalan Institution for Research and Advanced Studies (ICREA), Barcelona, Spain; Neuroscience Research Program, IMIM-Hospital Del Mar Research Institute, Barcelona, Spain

## Abstract

Maintaining appropriate circadian rhythmicity is essential for coordinating the activity of biological functions in mammalian organisms. A variety of physiological and behavioral changes have been associated with disturbances of this complex clock mechanism.

In the present study, we delve into the consequences of circadian arrhythmia using the Bmal1-knockout (KO) mouse line aiming to explore potential behavioral and motivational implications. We were able to identify the intricate activity patterns that define circadian disturbance in Bmal1-KO mice by utilizing a new analysis model based on entropy divergence.

Alterations in locomotor activity were accompanied by disruptions of circadian expression patterns in various clock genes as revealed by gene expression analysis. Additionally, we found a dysregulated gene expression profile in Bmal1-KO mice regarding genes related to circadian control in various brain nuclei. Specifically, the ventral striatum exhibited a dysregulation in the expression levels of genes modulating reward and motivation.

Further investigation revealed that BMAL1 deficient mice showed a sustained rise in motivation and seeking behavior for food and water reinforcers in the self-administration paradigm, independently of the caloric content of the reward. Together, our data reveal that disruptions in circadian rhythmicity, induced by alterations in the molecular clock, also impact the gene expression regulating the reward system. This, in turn, can lead to altered seeking behavior and motivation for natural rewards. In summary, the present study contributes to our understanding of how reward processing is under the regulation of circadian clock machinery.

## INTRODUCTION

The circadian clock controls the body’s natural rhythms by synchronizing them to the environment over a 24-hour period. Essential for maintaining internal temporal order, these molecular rhythms play a critical role in regulating a variety of biological processes, including sleep-wake cycles, hormone production and secretion, metabolic activities, and behavior (Fagiani et al., 2022; Kolbe et al., 2019; Serin & Acar Tek, 2019). The core architecture of the mammalian circadian clock relies on a complex molecular network featuring key components as the brain and muscle ARNT-like 1 (BMAL1) protein produced by the *Arntl* gene, and circadian locomotor output cycles kaput (*Clock*), or the paralog neuronal PAS domain protein 2 (*Npas2*). These proteins heterodimerize forming the CLOCK:BMAL1 or, alternatively, the NPAS2:BMAL1 complexes. These act as transcription factors binding to the E-box regions upstream thus promoting the transcription of the gene families Period and Cryptochrome (*Per* and *Cry*). These proteins will also heterodimerize into a PER:CRY complex that will inhibit the activity of the CLOCK:BMAL1 dimer and therefore, abolishing their own expression, establishing a negative transcriptional loop. Additional genes, such as *Rev-erb* and *Rora*, form a secondary self-regulation loop by repressing or enhancing *Bmal1* expression, respectively, which is known as the BMAL1 loop (Cox & Takahashi, 2019; Patton & Hastings, 2023). Moreover, the CLOCK:BMAL1 transcriptional factor binds to multiple E-boxes, controlling the transcription of numerous clock controlled-genes (CCGs) (Bozek et al., 2009). Therefore, regulating *Bmal1* expression is crucial for maintaining normal circadian rhythms in mammals. In this regard, the *Bmal1* knockout (Bmal1-KO) mouse line has emerged as an optimal animal model to study how arrythmia occurs in the circadian clock and how it can affect different systems (Rakai et al., 2014).

Notably, disruption of rhythmicity has also been linked to changes in motivation and reward modulation. At a molecular level, studies in both humans and mice have linked genetic modifications in *Bmal1*, *Per*, and *Clock* genes to increased drug use and altered reward processing (Castro-Zavala et al., 2022; Ciarleglio et al., 2008; Logan et al., 2014; Zhang et al., 2013). Alternatively, disruption of circadian rhythms can also be induced by environmental factors, many of which have long been described and researched in humans. For example, sleep deprivation, shift work, or jet-lag are associated to pathological states that impact reward and motivational characteristics as food and drug intake (DePoy et al., 2017; McClung, 2013).

In the realm of motivated behavior, the dopaminergic system plays a central role within the mesocorticolimbic circuit. This pivotal pathway extends from the ventral tegmental area (VTA) to the ventral striatum (vSTR), including the nucleus accumbens (NAc), and projections to the prefrontal cortex (PFC). Interestingly dopamine (DA) production and degradation are primarily regulated by the circadian cycle through the clock-controlled catabolic enzymes MAOA and MAOB, indicating a close interplay between these systems (Albrecht, 2017; Verwey et al., 2016). Furthermore, the dopaminergic system is subject to regulation by different neuromodulators, such as the endocannabinoid system, particularly through cannabinoid receptor 1 (CB1)-mediated actions (Parsons & Hurd, 2015).

The reward system responds to different types of cues in comparable and different ways (Pitchers et al., 2013). Food serves as a primary natural reward that acts as an essential stimulus for motivated responses and is also crucial for maintaining a proper metabolic state. Here, the hypothalamus, an integrated energy-sensing center, manages caloric and nutritional needs by sensing macronutrients and circulating regulatory hormones, neuropeptides, and neuromodulators like leptin, ghrelin, orexin/hypocretin, insulin, neuropeptide Y (NPY), and endocannabinoids (Barsh & Schwartz, 2002; Farr et al., 2016). Besides, this brain region contains the supraquiasmatic nucleus (SCN), the so-called central pacemaker (Cox & Takahashi, 2019). The SCN serves as a critical component that synchronizes metabolic regulation with daily changes in the environment by integrating information from circadian cues such as light or food intake. This integration allows for the coordination of appropriate behaviors in response to environmental stimuli. Together, the regulation of feeding behavior and energy homeostasis is a complex process that requires integration of a timekeeping system and feeding signals from the hypothalamus, but also motivational feeding signals from the mesolimbic pathways. Furthermore, the integration of homeostatic and reward-related feeding signals is facilitated by other higher-level decision-making centers, including the medial prefrontal cortex (mPFC), hippocampus, and amygdala, which are linked to the neurocircuitry of the motivation control (Baik, 2021; Liu et al., 2015).

While prior research has predominantly focused on investigating circadian disruption due to lack of *Bmal1* gene in the context of drugs of abuse such as cocaine (Castro-Zavala et al., 2022), this study aims to elucidate the impact of physiological arrhythmicity on homeostatic motivated behavior. To achieve this, we have utilized Bmal1-KO mice as an appropriate model for circadian arrhythmicity. Locomotor activity was analyzed employing a novel mathematical analysis method and leveraging a new software to assess rhythmicity using biological data. Additionally, we delved into the brain gene expression profile of these mice, focusing on the systems influencing the rewarding value of feeding behavior. Subsequently, we examined the motivational behavior of the animals through self-administration (SA) paradigms, using food and water as natural rewards. Our comprehensive approach unveiled the complex patterns that characterize circadian disruption consequences in behavior within our mouse model.

## MATERIALS AND METHODS

### Mice

Mice were obtained from the heterozygous C57BL/6 (Bmal1(+/-)) breeding colonies, kindly donated by Stem Cells and Cancer Lab at the IRB (Barcelona) and housed at UBIOMEX-PRBB. Animal rooms were maintained in a 12 h light/dark cycle (lights on at 7:30 am), with a constant temperature (21 ± 1°C) and humidity-(55% ± 10%). After weaning, mice were genotyped as homozygous Bmal1-KO (Bmal1(-/-)), heterozygous (Bmal1 (+/-)) or wild-type (WT) (Bmal1 (+/+)). Only Bmal1-KO and WT animals were used in the experiments. Both sexes males and females were used equally. All experiments were conducted in accordance with the guidelines of the European Communities Directive 88/609/EEC for animal research. Procedures were approved by the local ethical committee (CEEA-PRBB) and every effort was made to minimize animal suffering, discomfort and the number of animals used.

### Experimental design

We used different batches of animals as indicated for each experimental procedure. First, we analyzed the circadian rhythmicity of the animals through continuous and spontaneous locomotor activity measurements for 7 consecutive days (n=11/group). Time-series data was analyzed with a new mathematical analysis based on the variation of accumulated entropy (VAE).

A second group of Bmal1-KO and WT animals were used for the analysis of the rhythmicity of main clock genes *Per2*, *Cry2*, *Clock* and *Rev-erba* through a time-course gene expression analysis. Mice under *ad libitum* conditions (basal conditions) (n=3-4 per time-point and experimental group) were sacrificed every 4 h over a 24-h period (Zeitgeber time [ZT] 2, 6, 10, 14, 18 and 22, considering that ZT0 and ZT12 represent the onset of the light and dark phases, respectively) during which, samples from hypothalamus (HT) were collected for quantitative PCR (qPCR) experiments. Circadian rhythmicity of gene expression data was analyzed using Kronos, a computational tool to assess biological rhythms.

For the evaluation of differential gene expression between the WT and Bmal1-KO, custom OpenArrayTM platform was used. Animals were sacrificed (n=6 per group) at the same time-point (ZT18) and tissue corresponding to HT, mPFC and vSTR was collected. Then, the mRNA was extracted followed by RT-PCR and the resulting cDNA was analyzed in an OpenArrayTM plate (See Supp. Material S1).

In order to evaluate the motivational behavior of the animals, we exposed them to operant behavior paradigms where seeking and motivation for reinforcement was assessed. We conducted two different SA experiments using different animal batches. In the first SA (WT, n = 22; KO, n = 14), we used food as reinforcer while in the second one (WT, n = 14; KO, n = 14), water was used as a non-caloric reinforcer (Leib et al., 2016) to determine if the increase in reinforcement seeking could be affected by changes in metabolic balance resulting from the caloric component of the food. All the animals followed similar protocols with minor modifications and only food SA animals underwent the demand task as an extra behavioral evaluation test. All the SA sessions were performed throughout the dark phase of the light-dark cycle.

After the SA procedures, mice were sacrificed and tissue from HT and vSTR were dissected at ZT18 for molecular gene expression analysis with qPCR (WT, n = 15; KO, n = 9). We examined genes related to circadian rhythms and metabolic control within the HT (*AgRP, Npy, Avp, Vip* and *Hcrtr1)* and reward related control genes within the vSTR (*Drd1, Drd2, Maoa, Maob* and *Cnr1*).

### Spontaneous locomotor activity recording and analyses

For the assessment of spontaneous locomotion throughout the light/dark cycle (L:12, D:12), each mouse was individually housed in a cage (32 × 17 × 14 cm). Over seven consecutive days, the locomotor activity of each animal was recorded using the Panlab Infrared (IR) Actimeter (LE881 IR, Panlab s.l.u., Barcelona, Spain) and the SEDACOM software. The motor activity (MA) of a mouse was quantified as the number of counts per hour (MA/h).

### Entropy divergence analyses

For this analysis, we introduce a new analysis method called Variation of Accumulated Entropy (VAE). This method allows for determining the differences, in terms of information complexity, between two time series. Briefly (see Supplementary materials and methods for extended details), first, the Discrete Fourier Transform is applied to the different time series to obtain their *N* basic frequency components, i.e., harmonics, with their corresponding phases and amplitudes. Each of these harmonics contains a part of the total information contained in the original time series. Subsequently, a subset of the first *M* harmonics (*N*≥*M*≥0) is selected and the Inverse Discrete Fourier Transform is applied to this subset to obtain a partial reconstruction of the original time series. For each of these subsets of *M* harmonics, the Shannon information entropy (X. Liu et al., 2015) *E_M_* is calculated. In summary, a partial reconstruction of a time series using the first *M* harmonics of its decomposition in the Fourier spectrum will contain an informational complexity determined by *E_M_*. As the size of the subset of *M* harmonics increases, the reconstructed time series will be closer to the original and the contained information, determined by *E_M_*, will also be greater, i.e., the values of *E_M_* will increase as the size of the subset *M* used to reconstruct the time series increases.

For the time series obtained experimentally with the Bmal1-KO animals and WT, we define the VAE for a given subset of harmonics *M* as

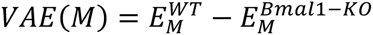

with 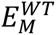 being the entropy associated with the subset formed by the first *M* harmonics of the time series obtained with the control animals, and 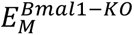 corresponding to that obtained with the Bmal1-KO mice. If beyond a certain critical value *M\**, the values of VAE stabilize, this indicates that considering more harmonics in the reconstruction of the time series, i.e., beyond a certain critical value (M*), the contributions to the accumulated entropy would not add significant information of the phenotypic characteristics of the mice that influences the obtained data series. This subset of *M\** harmonics contains the essential information that characterizes the circadian cycle in function of the specific characteristics of the animals. Consequently, applying the Inverse Discrete Fourier Transform to this subset is sufficient to reconstruct the time series of the circadian cycle determined by the specific phenotypic characteristics of the animals and discard other contributions to the time series, obtained experimentally, that depend on other factors. Furthermore, the value of VAE(M*) can be considered a metric that allows quantifying the distance, in terms of informational complexity contained in their circadian cycles, between the two sets of animals, WT and Bmal1-KO.

### Food self-administration procedure

Mice underwent food deprivation conditions for two days before the beginning of the self-administration procedure, maintaining the 85% of their initial weight as previously reported (Soria et al., 2005). Throughout the evaluation of food operant behavior, animals remained under the same food deprivation regime and were weighed after each session.

Mice were trained in the operant chambers (17.8 cm × 15.2 cm × 18.4 cm) (Med Associates, St. Albans, VT) to nose-poke for standard non-flavored food pellets (Noyes Precision Pellets, Research Diets Inc., USA) during 2-hour sessions. Nosepoking in the active hole of the chamber was rewarded with one food pellet and paired with a light cue. Inactive nosepokes yielded no output. For the first 10 days of the procedure, mice were trained under a fixed ratio (FR)1 schedule of reinforcement, where a nosepoke in the active hole resulted in the delivery of a single pellet. This was followed by 5 days under an FR3 schedule. Subsequently, a progressive ratio (PR) test was conducted, wherein the response requirement to earn a single reward escalated according to the following series: 1-2-3-5-12-18-27-40-60-90-135-200-300-450-675-1000. The breakpoint was defined as the number of nosepokes required to obtain the last pellet in the PR schedule. Finally, mice completed a demand task test modified from previous studies (Alegre-Zurano et al., 2022) where the criteria for earning a pellet followed an ascending trend every 10 minutes: 1, 2, 4, 6, 8, 11, 14, 18, 21, 28, 35, 42.

### Oral water self-administration procedure

Modified from (Soria et al., 2005), two days before beginning the experiment, water access was restricted to a single hour a day. This limitation aimed to increase the motivation for nosepoking and was maintained throughout the SA procedure. Again, animal’s body weight was daily monitored after each session. Food was available *ad libitum*. Mice were trained to receive water reinforcement under a fixed ratio (FR)1 (10 days) and FR3 (5 days) schedules of reinforcement. 24 h later, they underwent the PR test, mirroring the protocol used for food self-administration. The liquid dipper delivered water in 23 μl over 20 seconds, which were considered as a time-out period in which active nosepokes had no consequences. Each session finished either after 180 reinforcers were delivered or after 1 hour had passed.

### Tissue collection and RNA expression analyses

All the brain samples were collected using a 1mm brain matrix, placed in dry iced and immediately stored at -80°C. Total mRNA was extracted from brain tissue using TRIzol reagent, following the manufacturer’s instructions, and isopropanol for RNA precipitation. Then, cDNA was synthesized by reverse transcription of total mRNA. Time point gene expression and post behavioral molecular analysis were assessed through quantitative qPCR. The cDNA template was mixed with the primers (see Supp. Table 1) and SYBR Green PCR Master Mix. The expression level of each gene of interest within each sample was normalized against glyceraldehyde 3-phosphate dehydrogenase (*Gapdh*) and expressed relative to either WT group or to ZT2 of WT group for time-point gene expression. The fold change in gene expression of Bmal1-KO animals compared to controls was calculated using the 2-ΔΔCt method.

**Table 1.** Kronos circadian rhythmicity analysis data.

| Gene | Group | p.val | r.sq | avg | acro | amplitude | pairwise p-value |
| --- | --- | --- | --- | --- | --- | --- | --- |
| Per2 | WT | 0,15105 | 0,18042 | 0,92034 | 6,82234 | 0,30586 | 1 |
|  | KO | 0,52649 | 0,09398 | 1,23513 | 2,03947 | 0,13047 |  |
| Cry2 | WT | 0,01248 | 0,38556 | 0,78217 | 3,13735 | 0,15708 | 1 |
|  | KO | 0,08287 | 0,29937 | 1,04455 | 2,55370 | 0,15259 |  |
| Rev-erba | WT | 0,00087 | 0,54285 | 0,70612 | 4,23198 | 0,35255 | 0,002 |
|  | KO | 0,01798 | 0,46112 | 0,11940 | 4,31970 | 0,02056 |  |
| Clock | WT | 0,80448 | 0,02264 | 0,86396 | 2,80517 | 0,04304 | 1 |
|  | KO | 0,20947 | 0,21376 | 1,18056 | 3,04025 | 0,10491 |  |

### OpenArrayTM technology

To perform the OpenArray analyses, 2.5 μl of cDNA sample was combined with 2.5 μl TaqMan OpenArray Real-Time Master Mix (Thermo Fisher #4462159) and loaded into a single well of a 384-well plate (Abraham et al., 2021). Custom OpenArray plates were then automatically loaded using the AccuFill System (AccuFill System User Guide, PN4456986) and run in QuantStudio 12K. Amplification of the sequence of interest was normalized to reference endogenous genes, specifically, the geometric mean of *Actb*, *Gapdh*, *Hprt1*. Fold-change values were calculated using the ΔΔCt method. Data were analyzed with the ThermoFisher ConnectTM cloud web tool and Microsoft Excel.

### Statistical analysis

Normality (D’Agostino-Pearson and Kolmogorov-Smirnov tests), heteroscedasticity and homoscedasticity were assessed for all data sets. For single factor, two-group analyses involving parametric variables we employed unpaired Student’s *t-*tests. The study always considers the sex variable. If there are no differences, both sexes are analyzed as a single population. When an experimental condition followed a within-subject design a two-way ANOVA with repeated measures was conducted.

We harnessed Kronos, a computational tool by Bastiaanssen et al., (2023) to analyze biological rhythmicity efficiently in our study. This software specializes in evaluating circadian rhythms within biological datasets. Kronos proved instrumental in our study, streamlining rhythmicity analysis, and providing nuanced insights into oscillatory responses.

Demand curves were analyzed using the exponential model (Hursh & Silberberg, 2008): logQ = logQ_0_ + k (e−αQ_0_C − 1). In the exponential demand curve graphs, food pellet demand is plotted as a function of the price. Q_0_ represents consumption at a minimum price while α measures behavioral elasticity, assessesing demand responsiveness to commodity price shifts (Hursh, 1980). For the analysis of P_max_, we used the Excel calculator provided by Kaplan and collaborators (2014) (Kaplan & Reed, 2014). P_max_, known as the point of unit elasticity, is a metric used to evaluate motivation. The extra sum-of-squares F test was used to evaluate the fitness of the curves. This process was used to determine whether a single curve is sufficient to fit data from both WT and Bmal1-KO mice or whether distinct curves are more appropriate for the two datasets thereby indicating a distinct behavior between groups.

## RESULTS

### BMAL1 deficient mice show daily arrythmia in spontaneous locomotor activity and gene oscillation

To quantify arrhythmia status in locomotion or how one group differed from another by this parameter, we pioneered the use of Variation of Accumulated Entropy (VAE), based on the assumption that information in the corresponding time series of mice’s locomotor activity is altered as a result of disruptions to circadian rhythms. From the decomposition and mathematical processing of experimental data (See material and methods) VAE(M) is calculated. The results obtained show two distinct regions of behavior (Fig. 1a). In the first region (white area), VAE values changed significantly as the number of components (M) increases. This can be interpreted as each component of the subset M provides different information depending on whether it corresponds to the wild-type group or the Bmal1-KO animals. Therefore, this difference in the contained information is dependent on the specific characteristics of the analyzed experimental group. In the second region (colored area), starting from the value M*=110, it is observed that the VAE value exhibits significantly smaller variations, indicating that the new Fourier components added to reconstruct the curve of experimental data provide a similar amount of information to both time series. Therefore, they do not strongly rely on the attributes of each analyzed group and may relate to other shared factors. As a result, they do not significantly offer insights into circadian rhythms variation. Figures 1b and 1c display the original time series experimentally measured, S^WT^ (t) and S^Bmal1-KO^ (t) respectively. In these figures, the blue lines represent the reconstruction of the time series, applying the Inverse Fourier Transform, considering only the first M* Fourier components, which contain information dependent on the characteristics of each experimental group. The value of VAE M*=0.41 is a measure of the difference in the information complexity contained in the time series of locomotor activity. This positive value indicates that wild-type mice exhibit a more complex circadian dynamics regarding the amount of information contained in the time series compared to Bmal1-KO mice.

**Figure 1.**
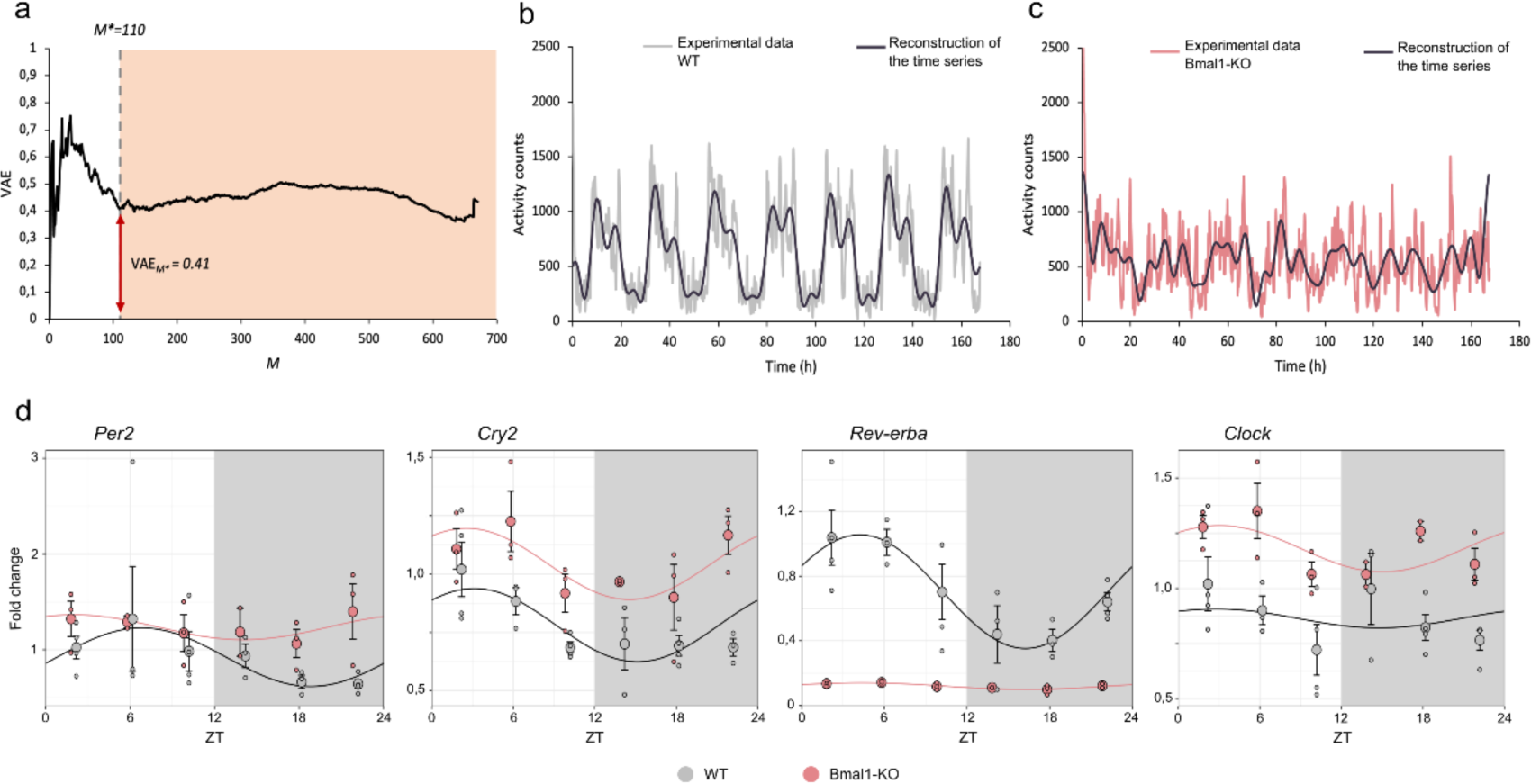
Assessment of daily arrhythmicity of Bmal1-KO mice. a. VAE values as a function of the number of Fourier components M used in their calculation. **b.** Reconstruction of the time series of locomotor activity in wild-type mice using only the first M*=110 Fourier components. **c.** Reconstruction of the time series of locomotor activity in Bmal-KO mice using only the first M*=110 Fourier components. (n = 11 mice per group) **d**. Graphical representation of time-course gene expression data of *Per2*, *Cry2*, *Rev-erba*, and *Clock* genes within the HT (n= 3 mice per group and time point). Graphs resulted from Kronos software show the cosine-fitted curves and SD from WT or Bmal1-KO mice. Period =24h, 12h light (white zone) 12h darkness (grey zone). ZT, Zeitgeber time, where ZT0 lights turn on.

The analysis of gene oscillation using the Kronos software facilitated a detailed analysis of rhythmicity within our experimental framework (see Supplementary materials and methods for the statistical details), leading to distinct observations regarding gene oscillations in WT and Bmal1-KO animals (Table 1). Figure 1d shows the regression curves delved by Kronos, fitted to the gene expression data along time, for both groups and for the examined clock genes. In WT animals, the *Cry2* gene exhibited significant oscillations (p < 0.05), indicating a pronounced cyclic pattern within the dataset. Conversely, in Bmal1-KO animals, the oscillations of the *Cry2* gene did not reach statistical significance (p = 0.082), suggesting a lack of observed rhythmicity. Despite the rhythmicity of *Rev-erba* gene for both genotypes, with p-values of 0.0008 (WT) and 0.018 (Bmal1-KO), the amplitude of this gene in the Bmal1-KO is close to 0, indicating low expression levels. However, pairwise analysis revealed differential rhythmicity between the curves (p < 0.01), implying distinct oscillatory patterns despite both groups showing significance in gene oscillation. Neither the *Per2* nor *Clock* genes exhibited significant oscillations in either WT or Bmal1-KO animals, suggesting an absence of discernible cyclic patterns in the expression of these genes within our dataset.

### Gene expression profile of the arrhythmic model

We evaluated the gene expression profile of genes related to relevant systems of our interest: clock genes, the reward system, and metabolic-related genes in three brain areas as HT, vSTR and mPFC. The genes that exhibited an absolute change greater than 1.3-fold in either direction with a p-value less than 0.05 revealed the differently expression for the Bmal1-KO group (Figure 2).

**Figure 2.**
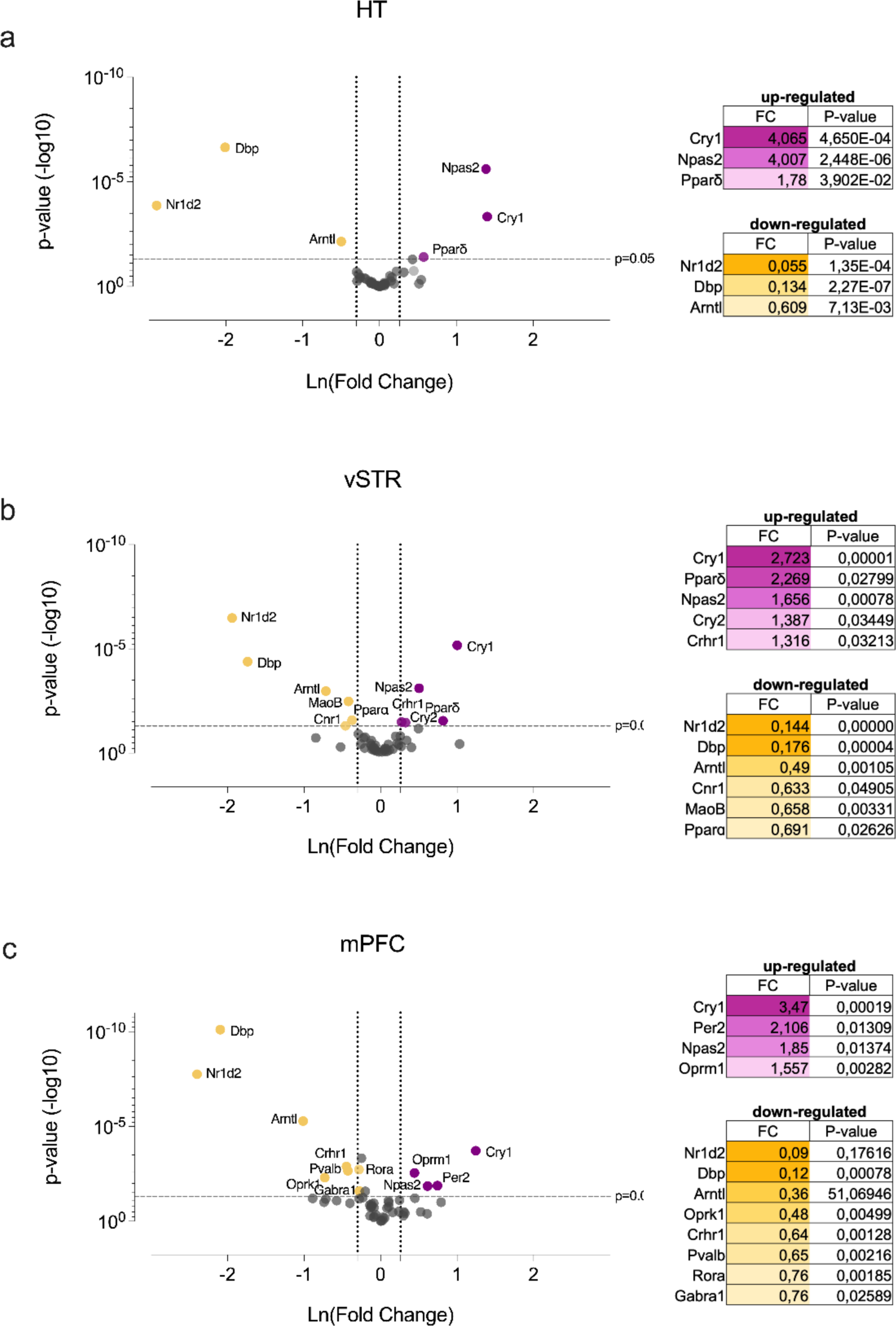
Differentially expressed genes of Bmal1-KO mice compared to WT mice. Comparisons of the expression of 52 genes assessed in OpenArray analysis of mRNA isolated from **a.** HT, **b.** vSTR and **c.** mPFC from WT mice (n = 6) and Bmal1-KO mice (n = 7). The volcano plot displays the relationship between fold change and significance between the two groups, applying a student’s t-test. The y-axis depicts the negative log10 of p-values of the t-tests (the horizontal slider corresponds to a p-value of 0.05) and the x-axis is the difference in expression between the two experimental groups as Ln fold changes (vertical sliders indicate mRNAs as either up-regulated (right area, purple dots) or down-regulated (left area, yellow dots) from a fold change (FC) of 1.3. The genes that exhibited both an absolute change greater than FC1.3 in either direction and a p-value <0.05 were exposed as the differently expressed genes.

Overall, in all nuclei examined, the clock molecular mechanism appears to be dysregulated. Clock genes were generally affected in every region: *Arntl*, *Dbp*, *Nr1d2* downregulated; *Npas2* and *Cry1* upregulated. Further, in HT we detected the upregulation of *Ppard* (Fig. 2a). Larger differences in the number of dysregulated genes were observed in the other two regions. Regarding the vSTR (Fig. 2b), the clock gene *Cry2*, was also upregulated. *Ppard* and *Crhr1* showed also significantly higher expression levels in the Bmal1-KO, while *Cnr1, Maob* and *Ppara* were downregulated. In the mPFC (Fig. 2c), we found dysregulation in other genes implicated in circadian rhythms machinery, *Rora* was downregulated and *Per2* was upregulated. However, genes related to other systems were also disrupted. *Oprm1* was upregulated whereas *Oprk1*, *Crhr1*, *Pvalb* and *Gabra1* showed a decreased expression.

### Circadian arrhythmicity alters food and water-induced self-administration behavior

We investigated the reward seeking alterations through SA paradigm reinforced with standard pellet of food, in which all mice successfully learned the task regardless of the reinforcement schedules (Figure 3b). The two-way ANOVA for the active nosepokes showed significant differences between groups with a much higher seeking behavior for the food reward in Bmal1-KO group (*Genotype effect* F(1, 32)= 37.96, p < 0.0001). The differences were maintained for a more demanding challenge within the test (FR3) (*Genotype effect* F(1, 32)= 43.53, p < 0.0001). No sex differences were found in neither FR schedules nor the PR (data not shown). The progressive ratio test revealed greater levels of motivation evidenced by a significant increase in active nosepokes (t32=3.318, p < 0.01) and a higher breaking point (t32=3.734, p < 0.001). Also, the Bmal1-KO mice obtained a much larger amount of food pellets overall (t20=2.872, p < 0.001).

**Figure 3.**
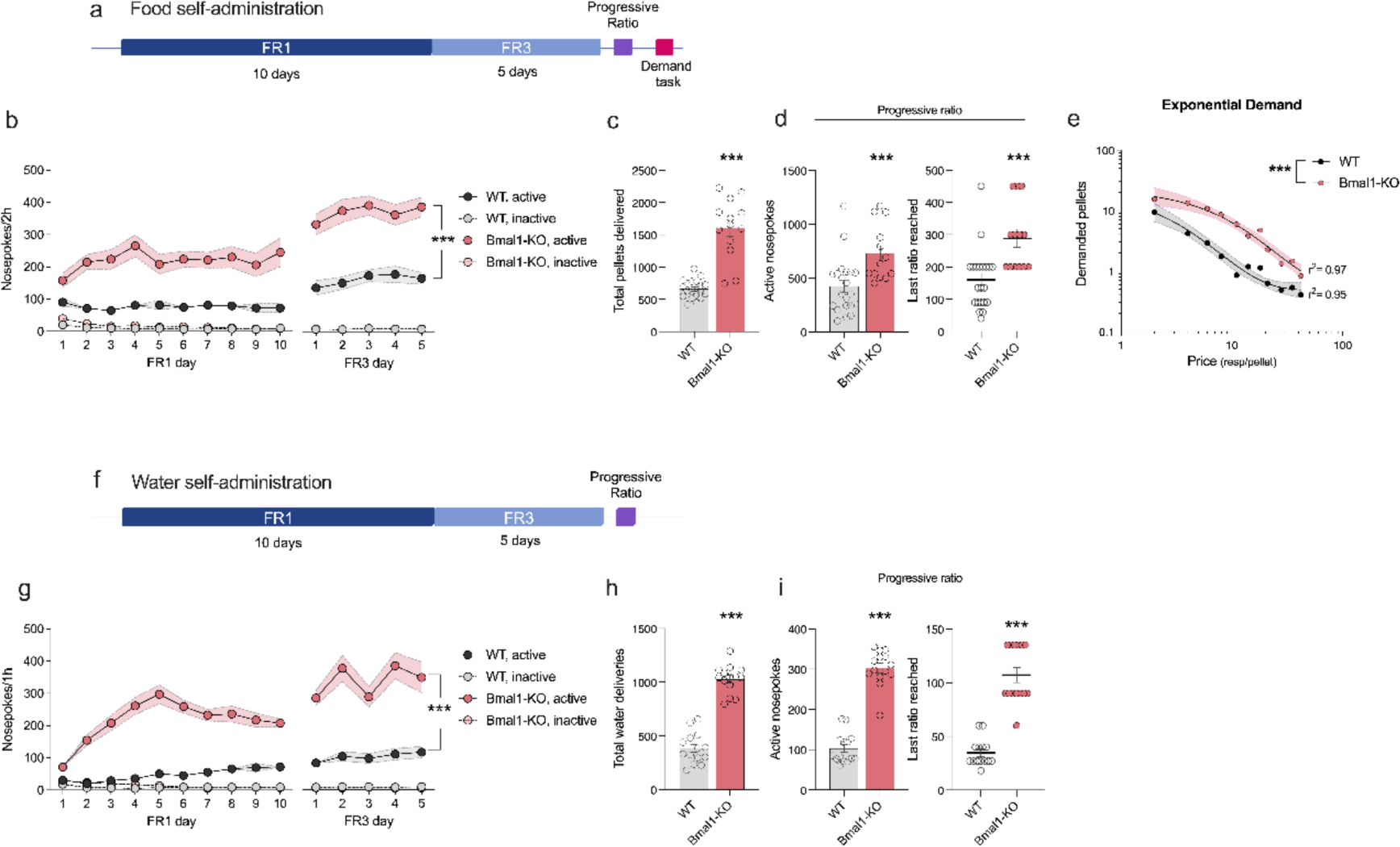
Mice operant training and motivation tests for caloric and non-caloric rewards. a. Schematic outline of the operant training behavior process for food rewards. **b**. Active and inactive nosepokes during FR1 and FR3 for food reward (WT n = 22; Bmal1-KO n = 14). **c.** Total pellets delivered through the 15 days of self-administration (SA) procedure. **d.** Total active nosepokes and breakpoint (last ratio reached) in the PR test for food. **e.** Behavioral economic analysis of the demand task and their exponential curve representation of Log of demanded pellets as a function of price. Extra sum-of-squares, ***p < 0.001. **f.** Schematic outline of the operant training behavior process for water rewards. **g**. Active and inactive nosepokes during FR1 and FR3 for water reward (WT n = 14; Bmal1-KO n = 14). **h.** Total water deliveries, obtained through the entire self-administration procedure. **I.** Total active nosepokes and breakpoint (last ratio reached) in the PR test in water SA. Two-way repeated measure ANOVA for each FR phase, ***p < 0.001. Student t-test, *p < 0.05; **p < 0.01; ***p < 0.001. See Supp. Table 2 for detailed statistical analysis.

Figure 3e illustrates the behavioral economics analysis of the demand task. Economic demand analysis facilitates understanding the role of effort in food procurement and the relationship between physiological and neural mechanisms (Rasmussen et al., 2016). The extra sum-of-squares F-test results showed that WT and Bmal1-KO mice exhibited different performance patterns during the task, best represented by two separate demand curves based on the Q0 and α parameters, (F(2, 17)= 84, p < 0.0001). Both groups exhibited a comparable preferred level of demanded pellets at a minimum price or Q0 (WT: 21,73, 95% CI [7.4, 36]; Bmal1-KO: 22,93, 95% CI [13, 33). However, Bmal1-KO mice displayed considerably less behavioral elasticity or α (WT: 0.0059, 95% CI [0.0033, 0.0084]; Bmal1-KO: 0.0018, 95% CI [0.0013, 0.0023]). The Pmax value for each data set was higher for Bmal1-KO mice (WT: 2.75; Bmal1-KO: 8,62), suggesting an increased motivation for the reward.

Following the same trend as in food self-administration, WT and Bmal1-KO mice exposed to water self-administration exhibited higher seeking behavior for water independently of the demanding conditions (*Genotype effect* FR1: F(1, 52)= 150.6, p < 0.0001; FR3:F(1, 52)= 61.43, p < 0.0001). Bmal1-KO mice also obtained a greater amount of water deliveries throughout the operant conditioning protocol (t26=12.36, p < 0.001; Figure 3h). Eventually, the Bmal1-KO animals presented an elevated number of active responses (t26=12.95, p <0.001) together with a significantly higher breakpoint in comparison with WT in the PR test (t26=9.298, p < 0.0001; Figure 3i).

### Dopaminergic system is altered in regions associated to reward control in Bmal1-KO mice

We considered to further examine the expression of several genes related to feeding behavior in the HT and reward, particularly to dopaminergic system within the vSTR.

Notably, we observed nearly identical alterations in gene expression within the HT in both at baseline and after exposure to food restriction period (Figure 4a,c) Specifically, there was a higher expression in the Bmal1-KO compared to WT in *AgRP* in both basal (t10=2.277, p < 0.05) and post food restriction (t20=2.872, p < 0.01) conditions. Additionally, there was a trend towards increased expression of *Npy* under basal conditions (t9=2.164, p = 0.0587), with an up-regulation observed after caloric restriction period (t20=2.589, p < 0.05). Further, *Avp* was significantly increased in the Bmal1-KO after the food self-administration (t21=3.790, p < 0.01) and showed a tendency towards this direction under basal conditions (t9=2.07, p = 0.068). Conversely, *Vip* displayed decreased expression in the Bmal1-KO group under both conditions (basal: t10=3.345, p < 0.01; post food restriction: t20=2.259, p < 0.05). Lastly, the gene for the hypocretin receptor 1 (*Hcrt1*) showed increased expression in Bmal1-KO animals compared to WT after food restriction (t21=2.406, p < 0.05) and displayed a similar trend under basal conditions (t10=2.196, p = 0.0528).

**Figure 4.**
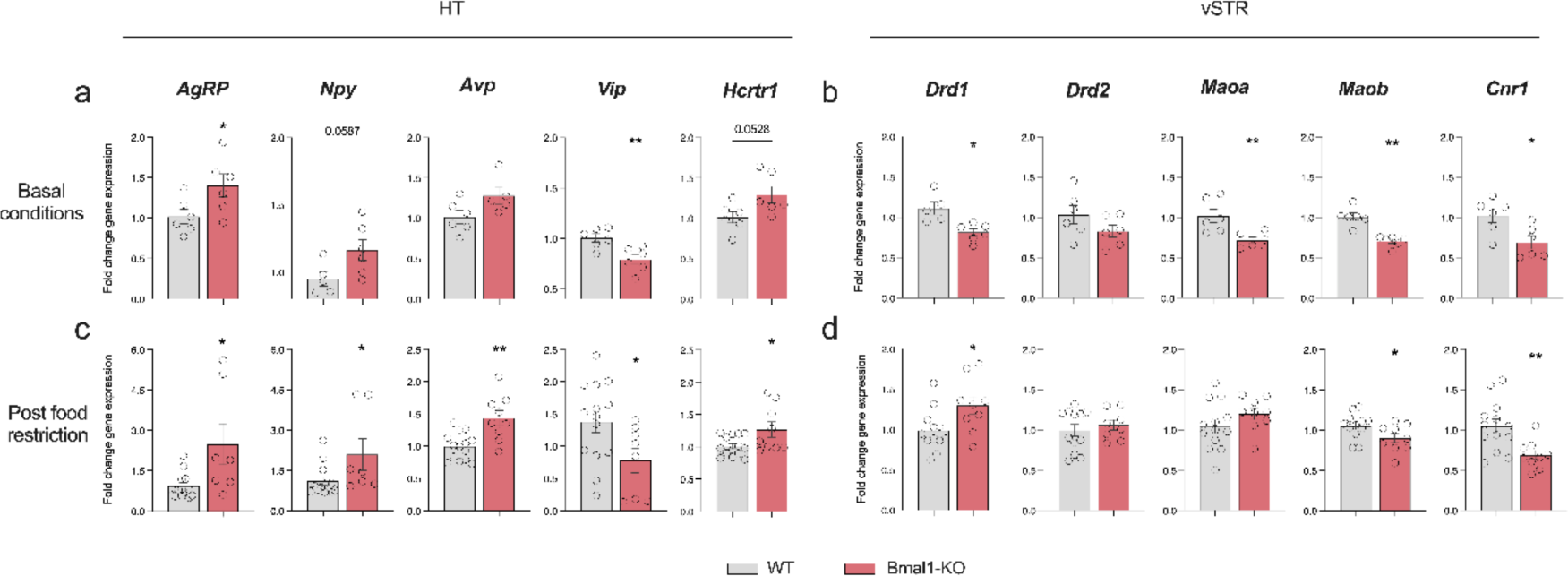
Bmal1-KO mice show altered expression of genes related to food intake control and the reward system in the HT and vSTR respectively. Gene expression analysis assessed by qPCR in basal conditions (WT n = 6; KO n = 6) in **a.** HT and **b.** vSTR; and after caloric restriction period (WT n = 15; KO n = 9) in **c.** HT and **d.** vSTR. Data are expressed as mean ± SEM. Student’s t-test (*p < 0.05; **p < 0.01; ***p < 0.001).

With respect to the vSTR (Figure 4b,d), we observed notable differences in expression between the different conditions, depending on the particular gene under study. In both conditions, *Drd2, Maob,* and *Cnr1* exhibited the same trend, while *Drd1* and *Maoa* did not. Caloric restriction seemed to upregulate *Drd1* expression in the Bmal1-KO group (t19=2.383, p < 0.05), while it was absent in the *ad libitum* condition (t9=3.436, p < 0.01). Alternatively, no differences were found in the expression of dopamine receptor 2 (*Ddr2*) in both conditions (basal: t10=1.489, ns; post food restriction: t18=0.6063, ns). Furthermore, *Maoa* and *Maob*, were down-regulated in Bmal1-KO mice in basal conditions (t10=3.301, p < 0.01; t10=5.509, p < 0.001). This downregulation was only mirrored in the food restriction condition for *Maob* (t20=2.109, p < 0.05). Finally, *Cnr1* displayed a downregulation in the Bmal1-KO mice compared to WT in both conditions (basal: t10=2.755, p < 0.05; post food restriction: t20=3.076, p < 0.01).

## DISCUSION

In this study, we establish that the absence of the *Bmal1* gene represents a model of circadian arrythmia in mice and how this alteration manifests in terms of activity due to the mutation. We assessed it through two different innovative techniques: for locomotor activity, the VAE analysis and time-course gene expression data analyzed with the computational tool Kronos. This arrythmia not only affects locomotion but also influences motivation and reward behavior of the animals. We further questioned whether this regulation is metabolic in nature or solely involves the reward system.

Our results offer additional evidence that the constitutive Bmal1-KO model used in this study displays a complex and less predictable activity patterns for several behaviors, including locomotion and motivated behaviors. We used the concept of variation of entropy to describe the degree of organization or randomness in the organism’s locomotor activity pattern. When the organism follows its normal circadian rhythm, the locomotor activity pattern displays a consistent and predictable structure. Therefore, peaks and troughs of the wave occur at approximately the same times every day, and the pattern will repeat in a regular and organized manner (Wu et al., 2018). Alternatively, disruption in the organism’s locomotor activity pattern caused by changes in the environment or a disruption in its internal clock, may result in a more random pattern. This means that peaks and troughs of the wave may occur at different times or in a less predictable manner, resulting in a higher degree of entropy and perturbation of the organism’s homeostasis. In agreement, Smith et al., (2023) showed impaired locomotor activity and feeding patterns in the same Bmal1-KO model, and Rakai et al., (2014) similarly demonstrated how another mouse model lacking BMAL1 struggled to adjust to light schedules and was completely arrhythmic under free-running conditions. Nevertheless, the novel mathematical method employed here allows the quantification of the degree of perturbation occurring in genetically modified mice, with VAE(M)=0.41 compared to the WT. In scenarios where an organism’s circadian rhythms may be disrupted by environmental, genetic, or pharmacological factors, this new measure can be used to quantify the extent of the disturbance and compare the impact of different situations on circadian rhythms.

Furthermore, chronobiology research has employed the fluctuation in the expression of clock genes during the day to ascertain the impact of genetic or environmental alterations on the circadian rhythms of animals at a molecular level (Girotti et al., 2009; Patel et al., 2016). Concretely, the study of the HT is of particular importance. It contains the SCN a nucleus that synchronizes the internal circadian timing system with the environment and regulates most circadian rhythms in the organism (Cox & Takahashi, 2019). Our investigation using Kronos (Bastiaanssen et al., 2023) unveiled distinctive gene-specific rhythmicity patterns in the HT within the studied animal groups. Within the HT, the *Cry2* gene showed significant oscillations exclusively in WT animals, whereas *Rev-erba* demonstrated significant oscillations in both WT and Bmal1-KO groups, albeit with significantly different rhythmic patterns probably due to the almost absence of expression of this gene in the Bmal1-KO. Conversely, *Per2* and *Clock* did not exhibit significant oscillatory behavior in either WT or Bmal1-KO animals. Since HT comprises different nuclei, the possible asynchrony between the rhythmic activity of different cellular groups could interfere the oscillatory behavior of *Per2* and *Clock*. However, these findings highlight the differential gene-specific rhythmicity present in WT and Bmal1-KO animals, shedding light onto the distinct oscillatory patterns of key genes and their potential implications in biological processes.

Consistently with the previous data, OpenArrayTM analysis reported a dysregulation in the expression of several clock genes in the three areas of study: HT, mPFC and vSTR. Said three areas were selected for their roles in the control of circadian rhythms and metabolic regulation, one of the key nuclei for reward modulation, and action control, respectively. Additionally, these latter two areas receive indirect projections from the SCN via the VTA and lateral habenula, (Becker-Krail et al., 2022), forming an intricate circuitry for regulating motivated behavior in a circadian-dependent manner.

Besides the impairment of the clock genes within the HT, as circadian machinery is directly or indirectly affected by *Bmal1* (Cox & Takahashi, 2019), the vSTR and mPFC also experienced up and down regulation of several clock genes which entails a dysregulation in the molecular circadian machinery. Particularly noteworthy is the general downregulation of the *Nr1d1* gene encoding REV-ERB, as expected since *Bmal1* directly regulates its expression (Preitner et al., 2002), possibly contributing to arrhythmicity and behavioral changes. All the alterations derived from the disturbances in this machinery may promote different behavioral alterations and pathological conditions as already reported in other studies (Porcu et al., 2020). For instance, resembling our results, Porcu and collegues (2020) found high expression of both *Cry1* and *Cry2* in NAc in mice that showed mood alterations. Conversely, the dysregulation in the expression of the other studied genes, may be a direct or indirect consequence of the disruption of the circadian molecular clocks. Specifically, overexpression of *Crhr1* may be a compensatory effect of the lower cortisol levels that are intrinsic in the Bmal1-KO line (Landgraf et al., 2014).

We intriguingly observed a sustained rise in motivation and seeking behavior in animals lacking BMAL1 in the SA paradigm, independent of the caloric content of the reward. This led us to further explore the possibility that the reward system is strongly affected by the lack of Bmal1, which induces rhythmicity impairments. Consequently, we investigated the dopaminergic system, one of the main systems implicated in reward regulation and influenced by circadian control (McClung, 2013). We found that dopaminergic signaling might be disturbed in the Bmal1-KO, as evidenced by a decrease in the expression of *Maob*, a DA-degrading enzyme. This coupled with the high seeking behavior displayed by Bmal1-KO mice in food SA procedures, is consistent with previous studies in which inhibition of the enzyme increased the effort required to obtain food rewards (Yohn et al., 2018). Additionally, we observed alterations in CB1 receptor gene, an important regulator of feeding and reward (Parsons & Hurd, 2015; Valverde et al., 2005; Wiley et al., 2005). The interplay of these systems could be contributing to the high motivation observed in these animals. Moreover, we reported dysregulation in *Gabra* and *Pvalb* genes, indicating a possible impairment of the GABAergic system within the mPFC. The dysregulation of this region can impact the ability to control impulses and regulate memory and decision making (Euston et al., 2012). Hence, it is possible that Bmal1-KO mice struggle with reduced self-control, which could result in an altered pursuit of primary rewards beyond basic survival needs, and therefore leading to compulsive-like behaviors. Supporting this hypothesis, our behavioral economic analysis revealed a low demand elasticity in the Bmal1-KO animals suggesting an impaired ability to adapt to changes in the price of reward and hinting a potential link between dysregulation of the molecular clock and a compromised behavioral adjustment in response to changing reward conditions. Furthermore, in the mPFC, the opioid system is disbalanced in the absence of BMAL1, possibly influencing the increase in motivation for natural rewards. Notably, we found, an upregulation of *Oprm1*, a mu receptor that enhances reward, and downregulation in *Oprk1* gene, a kappa receptor that promotes aversive behavior (Merrer et al., 2009) in mPFC.

Even though our research indicates that irregularities in circadian rhythmicity impact gene expression and, in turn, reward processing, it is crucial to recognize that different arrhythmicity models may yield different results. Contrary to our observations, some arrhythmicity models exhibit a decrease in motivation for food reward (Acosta et al., 2020). Similarly, in previous studies we reported a decrease in cocaine seeking of Bmal1-KO under a SA paradigm (Castro-Zavala et al., 2022). However, regulation of drug of abuse and natural rewards could be different and influenced by diverse environmental cues and internal conditions that could explain these apparent discrepancies. For instance, studies on constitutive Bmal1-KO mice (Welz et al., 2019) have revealed diverse phenotypic alterations, including accelerated ageing, reduced body weight, hypocortisol and hypoinsulinemia (Kondratov et al., 2006; Leliavski et al., 2014; Marcheva et al., 2010; Turek et al., 2005). In such a case, the metabolic issues may be impacting the motivation for cocaine and food in different ways. Further, regarding our results, the loss of BMAL1 leading to altered glucose (Kennaway et al., 2013) and ghrelin metabolism (Laermans et al., 2015) can incentive the food seeking by modulating significance of food as a mechanism aimed at maintaining metabolic homeostasis. Moreover, Bmal1-KO mice may be more vulnerable to stress conditions (Liu et al., 2015) which could lead to higher perception of the reinforcing value of food and a decrease in their impulsive control.

However, despite the metabolic impairments of Bmal1-KO model, the persistent increase in motivation beyond the caloric component of the reward, lead us to hypothesize that BMAL1 may be involved in influencing the reward system. Crucially, studies such as the one conducted by Kolbe et al., (2019), demonstrate that normalizing circadian rhythms can effectively normalize body weight and glucose metabolism, and could modulate the rewarding significance of food. Most of this emphasizes the importance of the lack of circadian rhythmicity as a key factor in the modulation of behavior.

Despite the potential impact of metabolic abnormalities resulting from disrupted circadian rhythms in mice, we contend that the primary driver behind the observed behaviors lies in the altered reward system. This assertion gains support from our consistent findings, even when utilizing a non-caloric reinforcement.

To explore whether arrhythmicity-induced metabolic changes contribute to the altered behavior in Bmal1-KO mice, we examined the expression of *AgRP* and *Npy*. Notably, their increased expression in Bmal1-KO, nearly independent of caloric restriction conditions, aligns with the activation of AgRP neurons. These neurons, co-expressing the orexigenic peptides AgRP and NPY, play an important role in feeding behavior and project to mesolimbic regions, regulating both reward-related behaviors and eating (Cansell et al., 2012). Our data also revealed an impact on the orexinergic (ORX) system, with increased *Hprtr1* expression in the hypothalamus, consistent with previous findings while indicating clock gene regulation of the ORX system (Feillet et al., 2017). Moreover, activation of D1 neurons mediates rewarding consumption and seeking, suggesting a pivotal role in the reinforcement process (Bond et al., 2020). Interestingly, the overexpression of D1 receptors within the vSTR after food SA, rather than in basal conditions, may be involved on excessive seeking behavior. This overexpression could perpetuate the hyperactivity of the same neuronal group, leading to the perseveration in seeking the reinforcer.

Recognizing the complexity of the interconnected roles of the circadian system, neurotransmitter control, metabolic homeostasis and reward processing, our comprehensive investigation provides insights into these intricate relationships.

In conclusion, our data illustrate that BMAL1 deficiency significantly disrupts circadian rhythms and the cellular molecular clock, which may affect gene expression and, consequently, the proper functioning of the reward system. This interplay results in changes in seeking behavior and motivation for natural rewards. While our findings provide insight into a specific aspect of this intricate relationship, the observed discrepancies emphasize the necessity of considering various models and their contextual factors. The broader implications highlight the importance of exploring both environmental and genetic origins of circadian disruptions to comprehensively understand their effects on reward processing. This research lays the foundation for further investigations delving into the dynamic interplay between circadian rhythms and the complex mechanisms governing the reward system.

## ABREVIATIONS

BMAL1: brain and muscle ARNT-like 1;
CB1: cannabinoid receptor 1;
Clock: circadian locomotor output cycles kaput;
Cry: Cryptochrome;
DA: Dopamine;
FR: fixed ratio;
HT: hypothalamus;
KO: Knockout;
mPFC: medial prefrontal cortex;
NPY: neuropeptide Y;
NAc: nucleus accumbens;
Per: Period;
PR: progressive ratio;
qPCR: quantitative PCR;
SA: self-administration;
SCN: supraquiasmatic nucleus;
VAE: variation of accumulated entropy;
vSTR: ventral striatum;
VTA: ventral tegmental area;
ZT: Zeitgeber time.

## AUTHOR CONTRIBUTIONS

P.B.-S., J.M., and O.V. were responsible for the study concept and design. P.B.-S., I.G.-L., J.M., L.A.-Z., A.C.Z.; OV carried out the experimental studies and performed the data analyses. P.S.W. and S.A.B. contributed to the data analysis while kindly providing the Bmal1 knockout mice strain. P.B.-S., I.G.-L., J.M. and O.V. drafted the manuscript. All authors critically reviewed the content and approved the final version for publication.

## REFERENCES

Abraham, N. A., Campbell, A. C., Hirst, W. D., & Nezich, C. L. (2021). Optimization of small-scale sample preparation for high-throughput OpenArray analysis. Journal of Biological Methods, 8(1), e143. 10.14440/jbm.2021.339

Acosta, J., Bussi, I. L., Esquivel, M., Höcht, C., Golombek, D. A., & Agostino, P. V. (2020). Circadian modulation of motivation in mice. Behavioural Brain Research, 382, 112471. 10.1016/j.bbr.2020.112471

Albrecht, U. (2017). Molecular Mechanisms in Mood Regulation Involving the Circadian Clock. Frontiers in Neurology, 8. https://www.frontiersin.org/articles/10.3389/fneur.2017.00030

Alegre-Zurano, L., Berbegal-Sáez, P., Luján, M. Á., Cantacorps, L., Martín-Sánchez, A., García-Baos, A., & Valverde, O. (2022). Cannabidiol decreases motivation for cocaine in a behavioral economics paradigm but does not prevent incubation of craving in mice. Biomedicine & Pharmacotherapy, 148, 112708. 10.1016/j.biopha.2022.112708

Baik, J.-H. (2021). Dopaminergic Control of the Feeding Circuit. Endocrinology and Metabolism, 36(2), 229–239. 10.3803/EnM.2021.979

Barsh, G. S., & Schwartz, M. W. (2002). Genetic approaches to studying energy balance: Perception and integration. Nature Reviews Genetics, 3(8), 589–600. 10.1038/nrg862

Bastiaanssen, T. F. S., Leigh, S.-J., Tofani, G. S. S., Gheorghe, C. E., Clarke, G., & Cryan, J. F. (2023). Kronos: A computational tool to facilitate biological rhythmicity analysis (p. 2023.04.21.537503). bioRxiv. 10.1101/2023.04.21.537503

Becker-Krail, D. D., Walker, W. H., & Nelson, R. J. (2022). The Ventral Tegmental Area and Nucleus Accumbens as Circadian Oscillators: Implications for Drug Abuse and Substance Use Disorders. Frontiers in Physiology, 13, 886704. 10.3389/fphys.2022.886704

Bond, C. W., Trinko, R., Foscue, E., Furman, K., Groman, S. M., Taylor, J. R., & DiLeone, R. J. (2020). Medial Nucleus Accumbens Projections to the Ventral Tegmental Area Control Food Consumption. The Journal of Neuroscience, 40(24), 4727–4738. 10.1523/JNEUROSCI.3054-18.2020

Bozek, K., Relógio, A., Kielbasa, S. M., Heine, M., Dame, C., Kramer, A., & Herzel, H. (2009). Regulation of Clock-Controlled Genes in Mammals. PLOS ONE, 4(3), e4882. 10.1371/journal.pone.0004882

Cansell, C., Denis, R., Joly-Amado, A., Castel, J., & Luquet, S. (2012). Arcuate AgRP neurons and the regulation of energy balance. Frontiers in Endocrinology, 3. https://www.frontiersin.org/articles/10.3389/fendo.2012.00169

Castro-Zavala, A., Alegre-Zurano, L., Cantacorps, L., Gallego-Landin, I., Welz, P.-S., Benitah, S. A., & Valverde, O. (2022). Bmal1-knockout mice exhibit reduced cocaine-seeking behaviour and cognitive impairments. Biomedicine & Pharmacotherapy, 153, 113333. 10.1016/j.biopha.2022.113333

Ciarleglio, C. M., Ryckman, K., Servick, S. V., Hida, A., Robbins, S., Wells, N., Hicks, J., Larson, S. A., Wiedermann, J. P., Carver, K., Hamilton, N., Kidd, K. K., Kidd, J. R., Smith, J. R., Friedlaender, J., McMahon, D. G., Williams, S., Summar, M. L., & Johnson, C. H. (2008). Genetic Differences in Human Circadian Clock Genes Among Worldwide Populations. Journal of Biological Rhythms, 23(4), 330–340. 10.1177/0748730408320284

Cox, K. H., & Takahashi, J. S. (2019). Circadian Clock Genes and the Transcriptional Architecture of the Clock Mechanism. Journal of Molecular Endocrinology, 63(4), R93–R102. 10.1530/JME-19-0153

DePoy, L. M., McClung, C. A., & Logan, R. W. (2017). Neural Mechanisms of Circadian Regulation of Natural and Drug Reward. Neural Plasticity, 2017, e5720842. 10.1155/2017/5720842

Euston, D. R., Gruber, A. J., & McNaughton, B. L. (2012). The Role of Medial Prefrontal Cortex in Memory and Decision Making. Neuron, 76(6), 1057–1070. 10.1016/j.neuron.2012.12.002

Fagiani, F., Di Marino, D., Romagnoli, A., Travelli, C., Voltan, D., Di Cesare Mannelli, L., Racchi, M., Govoni, S., & Lanni, C. (2022). Molecular regulations of circadian rhythm and implications for physiology and diseases. Signal Transduction and Targeted Therapy, 7(1). 10.1038/s41392-022-00899-y

Farr, O. M., Li, C. R., & Mantzoros, C. S. (2016). Central Nervous System Regulation of Eating: Insights from Human Brain Imaging. Metabolism: Clinical and Experimental, 65(5), 699–713. 10.1016/j.metabol.2016.02.002

Feillet, C. A., Bainier, C., Mateo, M., Blancas-Velázquez, A., Salaberry, N. L., Ripperger, J. A., Albrecht, U., & Mendoza, J. (2017). Rev-erbα modulates the hypothalamic orexinergic system to influence pleasurable feeding behaviour in mice. Addiction Biology, 22(2), 411–422. 10.1111/adb.12339

Girotti, M., Weinberg, M., & Spencer, R. (2009). Diurnal expression of functional and clock-related genes throughout the rat HPA axis: System-wide shifts in response to a restricted feeding schedule. American Journal of Physiology. Endocrinology and Metabolism, 296, E888–97. 10.1152/ajpendo.90946.2008

Hursh, S. R. (1980). Economic concepts for the analysis of behavior. Journal of the Experimental Analysis of Behavior, 34(2), 219–238. 10.1901/jeab.1980.34-219

Hursh, S. R., & Silberberg, A. (2008). Economic demand and essential value. Psychological Review, 115(1), 186–198. 10.1037/0033-295X.115.1.186

Kaplan, B. A., & Reed, D. D. (2014). Essential Value, Pmax, and Omax Automated Calculator. https://kuscholarworks.ku.edu/handle/1808/14934

Kennaway, D. J., Varcoe, T. J., Voultsios, A., & Boden, M. J. (2013). Global Loss of Bmal1 Expression Alters Adipose Tissue Hormones, Gene Expression and Glucose Metabolism. PLoS ONE, 8(6), e65255. 10.1371/journal.pone.0065255

Kolbe, I., Leinweber, B., Brandenburger, M., & Oster, H. (2019). Circadian clock network desynchrony promotes weight gain and alters glucose homeostasis in mice. Molecular Metabolism, 30, 140–151. 10.1016/j.molmet.2019.09.012

Kondratov, R. V., Kondratova, A. A., Gorbacheva, V. Y., Vykhovanets, O. V., & Antoch, M. P. (2006). Early aging and age-related pathologies in mice deficient in BMAL1, the core componentof the circadian clock. Genes & Development, 20(14), 1868–1873. 10.1101/gad.1432206

Laermans, J., Vancleef, L., Tack, J., & Depoortere, I. (2015). Role of the clock gene Bmal1 and the gastric ghrelin-secreting cell in the circadian regulation of the ghrelin-GOAT system. Scientific Reports, 5, 16748. 10.1038/srep16748

Landgraf, D., McCarthy, M. J., & Welsh, D. K. (2014). Circadian clock and stress interactions in the molecular biology of psychiatric disorders. Current Psychiatry Reports, 16(10), 483. 10.1007/s11920-014-0483-7

Leib, D. E., Zimmerman, C. A., & Knight, Z. A. (2016). Thirst. Current Biology, 26(24), R1260– R1265. 10.1016/j.cub.2016.11.019

Leliavski, A., Shostak, A., Husse, J., & Oster, H. (2014). Impaired Glucocorticoid Production and Response to Stress in Arntl-Deficient Male Mice. Endocrinology, 155(1), 133–142. 10.1210/en.2013-1531

Liu, J.-J., Mukherjee, D., Haritan, D., Ignatowska-Jankowska, B., Liu, J., Citri, A., & Pang, Z. P. (2015). High on food: The interaction between the neural circuits for feeding and for reward. Frontiers in Biology, 10(2), 165–176. 10.1007/s11515-015-1348-0

Logan, R. W., Williams III, W. P., & McClung, C. A. (2014). Circadian rhythms and addiction: Mechanistic insights and future directions. Behavioral Neuroscience, 128(3), 387–412. 10.1037/a0036268

Marcheva, B., Ramsey, K. M., Buhr, E. D., Kobayashi, Y., Su, H., Ko, C. H., Ivanova, G., Omura, C., Mo, S., Vitaterna, M. H., Lopez, J. P., Philipson, L. H., Bradfield, C. A., Crosby, S. D., JeBailey, L., Wang, X., Takahashi, J. S., & Bass, J. (2010). Disruption of the clock components CLOCK and BMAL1 leads to hypoinsulinaemia and diabetes. Nature, 466(7306), 627–631. 10.1038/nature09253

McClung, C. A. (2013). How Might Circadian Rhythms Control Mood? Let Me Count the Ways... Biological Psychiatry, 74(4), 242–249. 10.1016/j.biopsych.2013.02.019

Merrer, J. L., Becker, J. A. J., Befort, K., & Kieffer, B. L. (2009). Reward Processing by the Opioid System in the Brain. Physiological Reviews, 89(4), 1379–1412. 10.1152/physrev.00005.2009

Parsons, L. H., & Hurd, Y. L. (2015). Endocannabinoid signaling in reward and addiction. Nature Reviews. Neuroscience, 16(10), 579–594. 10.1038/nrn4004

Patel, S. A., Velingkaar, N., Makwana, K., Chaudhari, A., & Kondratov, R. (2016). Calorie restriction regulates circadian clock gene expression through BMAL1 dependent and independent mechanisms. Scientific Reports, 6(1), 25970. 10.1038/srep25970

Patton, A. P., & Hastings, M. H. (2023). The Mammalian Circadian Time-Keeping System. *Journal of Huntington’s Disease*, Preprint(Preprint), 1–14. 10.3233/JHD-230571

Pitchers, K. K., Vialou, V., Nestler, E. J., Laviolette, S. R., Lehman, M. N., & Coolen, L. M. (2013). Natural and Drug Rewards Act on Common Neural Plasticity Mechanisms with ΔFosB as a Key Mediator. The Journal of Neuroscience, 33(8), 3434–3442. 10.1523/JNEUROSCI.4881-12.2013

Porcu, A., Vaughan, M., Nilsson, A., Arimoto, N., Lamia, K., & Welsh, D. K. (2020). Vulnerability to helpless behavior is regulated by the circadian clock component CRYPTOCHROME in the mouse nucleus accumbens. Proceedings of the National Academy of Sciences, 117(24), 13771–13782. 10.1073/pnas.2000258117

Preitner, N., Damiola, F., Luis-Lopez-Molina, Zakany, J., Duboule, D., Albrecht, U., & Schibler, U. (2002). The Orphan Nuclear Receptor REV-ERBα Controls Circadian Transcription within the Positive Limb of the Mammalian Circadian Oscillator. Cell, 110(2), 251–260. 10.1016/S0092-8674(02)00825-5

Rakai, B. D., Chrusch, M. J., Spanswick, S. C., Dyck, R. H., & Antle, M. C. (2014). Survival of adult generated hippocampal neurons is altered in circadian arrhythmic mice. PloS One, 9(6), e99527. 10.1371/journal.pone.0099527

Rasmussen, E. B., Robertson, S. H., & Rodriguez, L. R. (2016). The Utility of Behavioral Economics in Expanding the Free-Feed Model of Obesity. Behavioural Processes, 127, 25–34. 10.1016/j.beproc.2016.02.014

Serin, Y., & Acar Tek, N. (2019). Effect of Circadian Rhythm on Metabolic Processes and the Regulation of Energy Balance. Annals of Nutrition and Metabolism, 74(4), 322–330. 10.1159/000500071

Smith, J. G., Koronowski, K. B., Mortimer, T., Sato, T., Greco, C. M., Petrus, P., Verlande, A., Chen, S., Samad, M., Deyneka, E., Mathur, L., Blazev, R., Molendijk, J., Kumar, A., Deryagin, O., Vaca-Dempere, M., Sica, V., Liu, P., Orlando, V., Parker, B. L., Baldi, P., Welz, P.-S., Jang, C., Marsi, S., Benitah, S. A., Muñoz-Cánoves, P., & Sassone-Corsi, P. (2023). Liver and muscle circadian clocks cooperate to support glucose tolerance in mice. Cell Reports, 42(6), 112588. 10.1016/j.celrep.2023.112588

Soria, G., Mendizábal, V., Touriño, C., Robledo, P., Ledent, C., Parmentier, M., Maldonado, R., & Valverde, O. (2005). Lack of CB1 cannabinoid receptor impairs cocaine self-administration. Neuropsychopharmacology: Official Publication of the American College of Neuropsychopharmacology, 30(9), 1670–1680. 10.1038/sj.npp.1300707

Turek, F. W., Joshu, C., Kohsaka, A., Lin, E., Ivanova, G., McDearmon, E., Laposky, A., Losee-Olson, S., Easton, A., Jensen, D. R., Eckel, R. H., Takahashi, J. S., & Bass, J. (2005). Obesity and metabolic syndrome in circadian Clock mutant mice. Science, 308(5724), 1043–1045. 10.1126/science.1108750

Valverde, O., Karsak, M., & Zimmer, A. (2005). Analysis of the endocannabinoid system by using CB1 cannabinoid receptor knockout mice. Handbook of Experimental Pharmacology, 168, 117–145. 10.1007/3-540-26573-2_4

Verwey, M., Dhir, S., & Amir, S. (2016). Circadian influences on dopamine circuits of the brain: Regulation of striatal rhythms of clock gene expression and implications for psychopathology and disease. F1000Research, 5, F1000 Faculty Rev-2062. 10.12688/f1000research.9180.1

Welz, P.-S., Zinna, V. M., Symeonidi, A., Koronowski, K. B., Kinouchi, K., Smith, J. G., Guillén, I. M., Castellanos, A., Furrow, S., Aragón, F., Crainiciuc, G., Prats, N., Caballero, J. M., Hidalgo, A., Sassone-Corsi, P., & Benitah, S. A. (2019). BMAL1-Driven Tissue Clocks Respond Independently to Light to Maintain Homeostasis. Cell, 177(6), 1436–1447.e12. 10.1016/j.cell.2019.05.009

Wiley, J. L., Burston, J. J., Leggett, D. C., Alekseeva, O. O., Razdan, R. K., Mahadevan, A., & Martin, B. R. (2005). CB1 cannabinoid receptor-mediated modulation of food intake in mice. British Journal of Pharmacology, 145(3), 293–300. 10.1038/sj.bjp.0706157

Wu, M., Zhou, F., Cao, X., Yang, J., Bai, Y., Yan, X., Cao, J., & Qi, J. (2018). Abnormal circadian locomotor rhythms and Per gene expression in six-month-old triple transgenic mice model of Alzheimer’s disease. Neuroscience Letters, 676, 13–18. 10.1016/j.neulet.2018.04.008

Yohn, S. E., Reynolds, S., Tripodi, G., Correa, M., & Salamone, J. D. (2018). The monoamine-oxidase B inhibitor deprenyl increases selection of high-effort activity in rats tested on a progressive ratio/chow feeding choice procedure: Implications for treating motivational dysfunctions. Behavioural Brain Research, 342, 27–34. 10.1016/j.bbr.2017.12.039

Zhang, L., Ptáček, L. J., & Fu, Y.-H. (2013). Diversity of Human Clock Genotypes and Consequences. Progress in Molecular Biology and Translational Science, 119, 51–81. 10.1016/B978-0-12-396971-2.00003-8

