## Supplementary material for "Lack of rhythmicity in Bmal1 deficient mice impairs motivation towards natural stimuli": SUPPLEMENTARY MATERIALS AND METHODS.pdf

#### Variation of accumulated entropy (VAE)

To compare both series, the Discrete Fourier Transform is applied to obtain a decomposition of each time series into a set of N elementary components (Supp. Fig. 1), each of which corresponds to a specific frequency and its respective amplitude and phase, *i.e.*:

$$S(t) = \sum_{i=0}^{N} A_i \cdot f(\omega_i, \varphi_i)$$

Here S(t) represents the time series data, N is the number of components (harmonics), and f(ω_i,φ_i) is a sinusoidal function of frequency ωi and phase φi. Finally, Ai is the amplitude with which the i-th component contributes to the time series S(t). When the Inverse Fourier Transform is applied to a subset M of these Fourier components (with M<N), a partial reconstruction Sp(t) of the original series S(t) is obtained, *i.e.*:

$$S_p(t) = \sum_{i=0}^{M} A_i \cdot f(\omega_i, \varphi_i)$$

As a result Sp(t) only contains a part of the total information within the S(t). The amount of information contained in Sp(t) can be described by calculating the Shannon entropy (X. Liu et al., 2015) of the sub-series formed by M Fourier components. To do this, the normalized distribution of the different components was calculated according to:

$$p_i = \frac{A_i}{\sum_{j=0}^{M} A_j}$$

and the corresponding entropy of the sub-series

$$E_M = -\sum_{i=0}^{M} p_i \cdot log\,(p_i)$$

According to the described method, this entropy of sub-series M can be calculated with the temporal data of the Bmal1-KO and WT mice, i.e., $E_M$^WT. Finally, the variation of accumulated entropy (VAE) is calculated as the sum of the entropy differences between the experimental group of animals and control (WT) animals for each of the possible sub-series of size M

$$\Delta E_M = \sum_{j=0}^{M} (E_M^{WT} - E_M^{P})$$

According to this measure, as long as the sub-series of size M contains information about circadian cycles, which are dependent on the specific characteristics of the analyzed animals, the values of $\Delta E_M$ will change as the size of the sub-series M increases. Conversely, beyond a certain critical value M*, the contributions to the accumulated entropy from the added Fourier components will contain information associated with other non-intrinsic aspects of the analyzed animals but rather external factors, i.e., environmental noise. This will result in the value of $\Delta E_M$ remaining stable as the size M of the sub-series increases, and therefore, these new components do not provide additional significant information about the characteristics of the mice's circadian rhythms. The value of VAE calculated at the point M*, i.e., ΔE_(M^*), can be considered a measure of the difference in the amount of information associated with circadian cycles contained in the time series. The sign of ΔE_(M^*) indicates which of the series contains more information, i.e., ΔE_(M^*)>0 indicates that the time series of control animals are more complex in terms of information than the corresponding series of experimental animals, and vice versa in the case of ΔE_(M^*)<0.

**Kronos**

Kronos predicts sinusoid curves for each variable based on a specified period, generating a kronosOut object with valuable outputs: variance proportion explained by predicted curves, corresponding p-values, acrophase, and amplitude. It also offers model details beneficial for prediction and statistical applications.

To compare rhythmicities between groups, Kronos employs a generalized linear model on user-defined categorical predictors, decomposed sine and cosine components, and interactions. The resulting kronosOut object encompasses interaction details, aggregating factor-related p-values with Bonferroni's adjustment for the lowest adjusted p-value. Additionally, it facilitates pairwise comparisons, distinguishing overall differences from group-specific rhythmicity variations. Adhering to Kronos' original methods, we estimated rhythm characteristics, focusing on vital 24-hour cycles crucial for maintaining health and synchronizing physiological processes. Differential rhythmicity analysis unveiled oscillatory signal responses to internal and external factors, contributing to significant biological insights.

**Weigh control**

We weighed the animals daily throughout both self-administration procedures to control the effects of food and water restriction (Supp. Fig. 2). The weight changes for both groups were different in the early days (Supp. Fig. 2b,e) but it stabilized along the procedure, and we couldn't see any significant difference in weight changes between the two groups by the end of the protocol (Supp. Fig. 2c,f). See Supp. Table 3 for statistical analyses.

### Supplementary Table 1. Primer sequences used for qPCR.

| ID | Sequence FWD | Sequence REV |
|---|---|---|
| *Gapdh* | 5'- GGA GAA ACC TGC CAA GTA TGA -3' | 5'- TCC TCA GTG TAG CCC AAG A -3' |
| *Clock* | 5'- GGT CAA GGG CTA CAG ATG TTT -3' | 5'- CAG GTG TGA GTT GCT GGA TAT TA -3' |
| *Per2* | 5'- CAA CAA CCC ACA CAC CAA AC -3' | 5'- CTC GAT CAG ATC CTG AGG TAG A -3' |
| *Cry2* | 5'- GAG AAC CAT GAC GAC ACC TAT G -3' | 5'- AGC TTC TGT CTC TCC TCC TT -3' |
| *Nr1d1* | 5'- CTT CAT CCT CCT CCT CCT TCT A -3' | 5'- GTA ATG TTG CTT GTG CCC TTG -3' |
| *Vip* | 5'- CCG TCT TCA CAG ATA ACT ACA C -3' | 5'- GCC TCT TCC CAT CAT TTC TC -3' |
| *Avp* | 5'- GCT CGC CAG GAT GCT CAA -3' | 5'- ACA CTG TCT CAG CTC CAT GTC AGA -3' |
| *Npy* | 5'- CTC ATC ACC AGA CAG AGA TAT G -3' | 5'- GGT CTT CAA GCC TTG TTC T -3' |
| *Drd1* | 5'- CTC CAT CTC CAA GGA CTG TAA TC -3' | 5'- CTT CTC CAG TGG CTT AGG TAT G -3' |
| *Drd2* | 5'- CAC AGA CCA GAA TGA GTG TAT C -3' | 5'- CAT CCT TGA GTG GTG TCT TC -3' |
| *AgRp* | 5'- AGG TGC TAG ATC CAC AGA A -3' | 5'- ATT GAA GAA GCG GCA GTA G -3' |
| *Hcrtr1* | 5'- CCT TAA GAG AGT GTT CGG GAT G -3' | 5'- TTG CCA CTG AGG AAG TTG TAG -3' |
| *Cnr1* | 5'- AGG AGA CAC AAC CAA CAT TAC A -3' | 5'- TGA AGC ACT CCA TGT CCA TAA A -3' |
| *Maoa* | 5'- GGG CGG TAC AAG GGT CTG TT -3' | 5'- ACG TCG AAC ATG TGG CCT GT -3' |
| *Maob* | 5'- GTG GAC CTT GGA GGA TCT TAT G -3' | 5'- CAG CCG CTC AAC TTC ATT AAC -3' |

**Supplementary Table 2. Statistical table related to Figure 3.**

| Figure | Test | Source of variation | Test value | p-value | Significant |
|---|---|---|---|---|---|
| Figure 3b - FR1 | Two-way ANOVA | Time x Genotype | F (9, 288) = 3,559 | P=0,0003 | Yes |
| Figure 3b - FR1 | Two-way ANOVA | Time | F (1,804, 57,73) = 2,357 | P=0,1088 | No |
| Figure 3b - FR1 | Two-way ANOVA | Genotype | F (1, 32) = 36,82 | P<0,0001 | Yes |
| Figure 3b - FR3 | Two-way ANOVA | Time x Genotype | F (4, 128) = 0,7007 | P=0,5928 | No |
| Figure 3b - FR3 | Two-way ANOVA | Time | F (2,379, 76,12) = 3,227 | P=0,0371 | Yes |
| Figure 3b - FR3 | Two-way ANOVA | Genotype | F (1, 32) = 43,08 | P<0,0001 | Yes |
| Figure 3g - FR1 | Two-way ANOVA | Time x Genotype | F (9, 234) = 9,152 | P<0,0001 | Yes |
| Figure 3g - FR1 | Two-way ANOVA | Time | F (3,604, 93,70) = 14,08 | P<0,0001 | Yes |
| Figure 3g - FR1 | Two-way ANOVA | Genotype | F (1, 26) = 145,1 | P<0,0001 | Yes |
| Figure 3g - FR3 | Two-way ANOVA | Time x Genotype | F (4, 104) = 1,380 | P=0,2461 | No |
| Figure 3g - FR3 | Two-way ANOVA | Time | F (3,205, 83,34) = 3,251 | P=0,0233 | Yes |
| Figure 3g - FR3 | Two-way ANOVA | Genotype | F (1, 26) = 63,72 | P<0,0001 | Yes |

**Supplementary Table 3. Statistical table related to Supplementary Figure 2.**

| Figure | Test | Source of variation | Test value | p-value | Significant |
|---|---|---|---|---|---|
| Figure S2b | Two-way ANOVA | Interaction | F (1, 29) = 0,3585 | P=0,5540 | No |
| Figure S2b | Two-way ANOVA | Genotype | F (1, 29) = 25,90 | P<0,0001 | Yes |
| Figure S2b | Two-way ANOVA | Sex | F (1, 29) = 0,6604 | P=0,4230 | No |
| Figure S2c | Two-way ANOVA | Interaction | F (1, 30) = 0,7393 | P=0,3967 | No |
| Figure S2c | Two-way ANOVA | Genotype | F (1, 30) = 1,998 | P=0,1678 | No |
| Figure S2c | Two-way ANOVA | Sex | F (1, 30) = 1,528 | P=0,2260 | No |
| Figure S2e | Two-way ANOVA | Interaction | F (1, 24) = 0,2880 | P=0,5964 | No |
| Figure S2e | Two-way ANOVA | Genotype | F (1, 24) = 6,499 | P=0,0176 | Yes |
| Figure S2e | Two-way ANOVA | Sex | F (1, 24) = 2,450 | P=0,1306 | No |
| Figure S2f | Two-way ANOVA | Interaction | F (1, 22) = 0,005117 | P=0,9436 | No |
| Figure S2f | Two-way ANOVA | Genotype | F (1, 22) = 3,529 | P=0,0736 | Yes |
| Figure S2f | Two-way ANOVA | Sex | F (1, 22) = 9,648 | P=0,0052 | Yes |

**Supplementary Fig. 1. a**. Fourier spectrum of the time series of locomotor activity in Wild-Type mice. **b.** Fourier spectrum of the time series of locomotor activity in Bmal1-KO mice.

**Supplementary Fig. 2. Changes in weight of mice along the food and water restriction conditions. a.** Weight of the animals along the self-administration days under food and **d.** water restriction conditions. Change in weight was different for both genotypes after 5 days of **b.** food or **e.** water deprivation. Change in weight was equal for both mice groups by the end of the procedure (day 15) maintaining **c.** food or **f.** water restriction conditions. Two-way ANOVA, $p < 0.05$; $^{**}p < 0.01$; $^{***}p < 0.001$.
