## Supplementary figures and images for "Lack of rhythmicity in Bmal1 deficient mice impairs motivation towards natural stimuli"

### Supp. Fig. 1.tiff

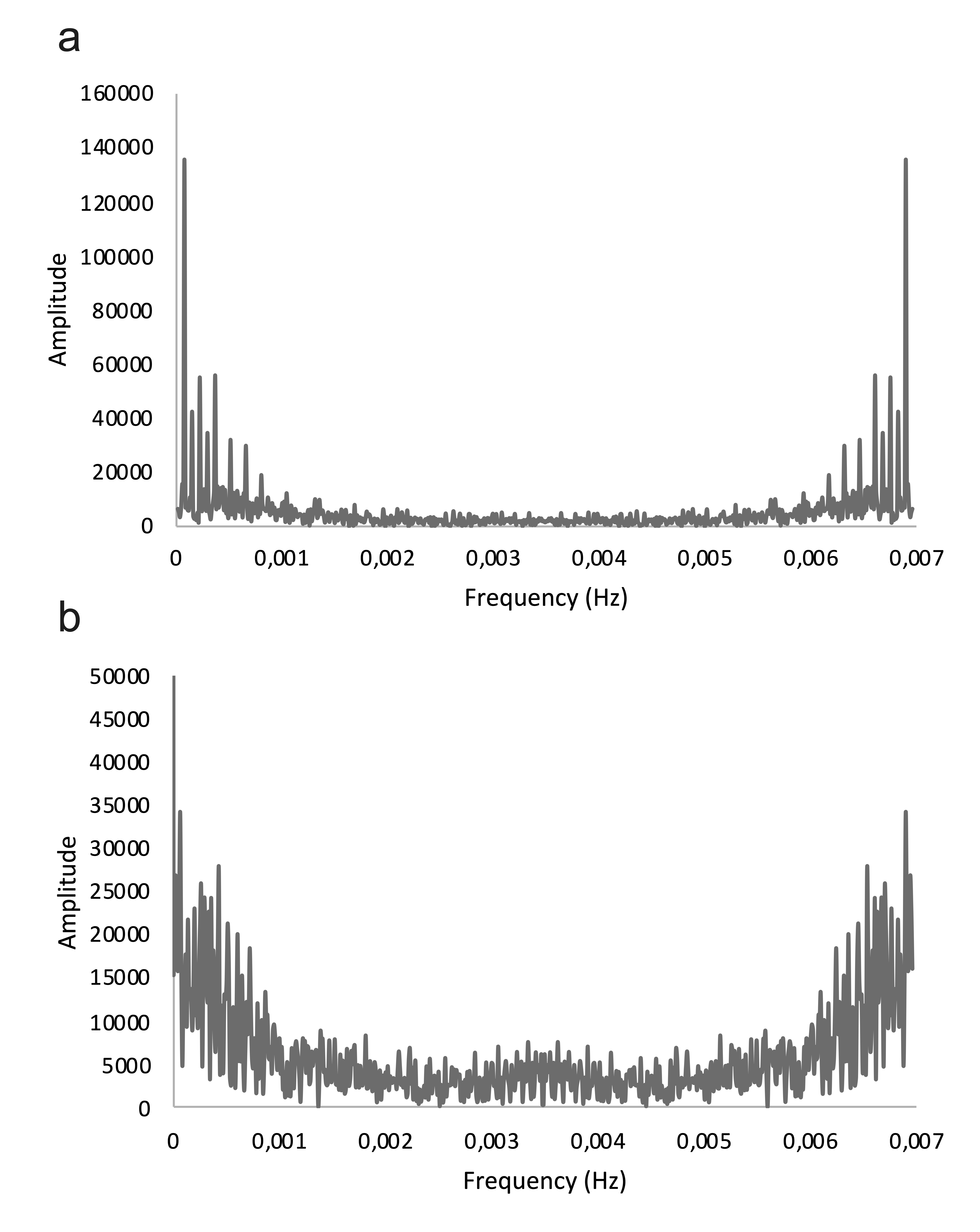

### Supp. Fig. 2.tiff

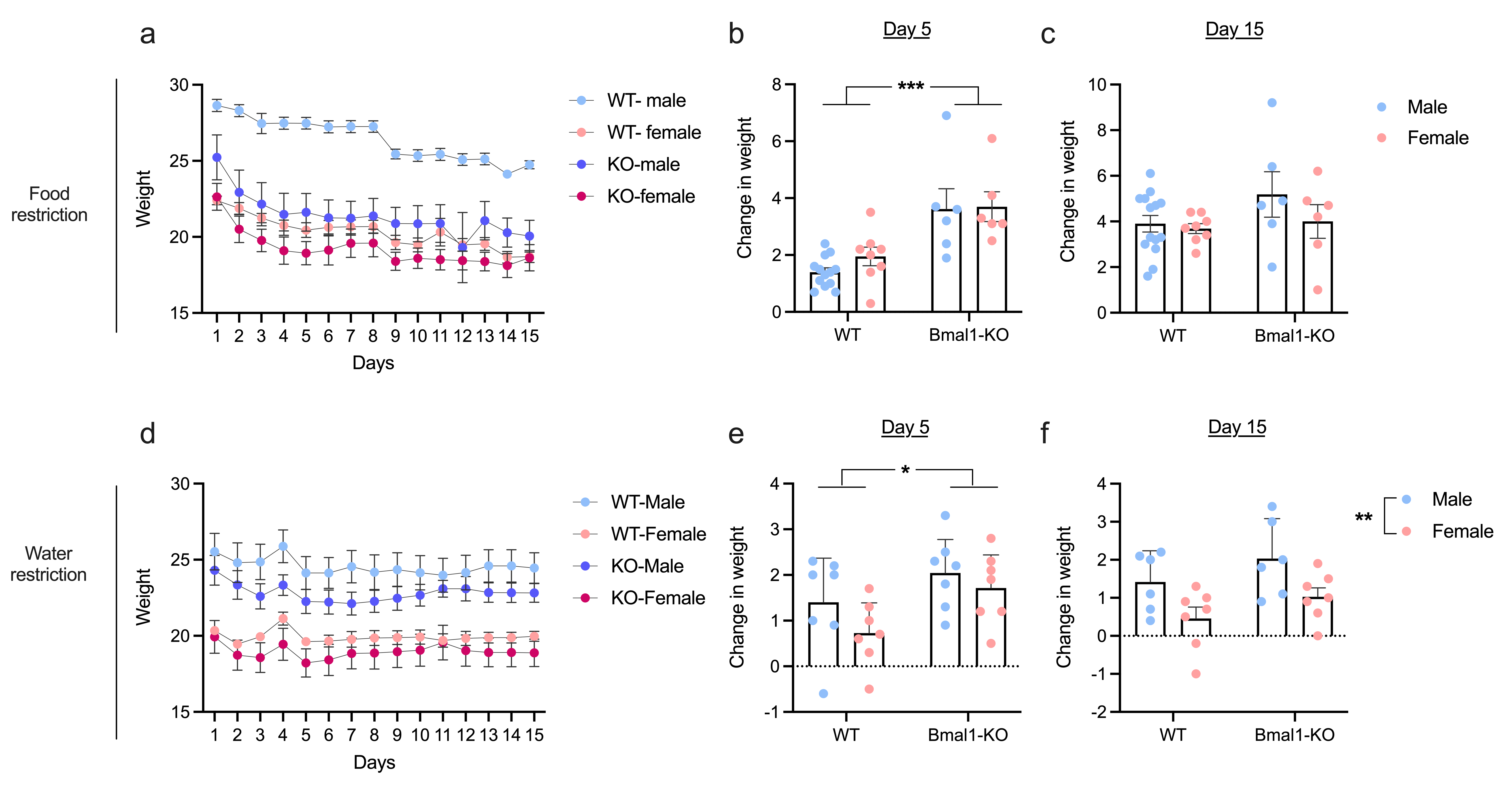
